# Information Isometry Technique Reveals Organizational Features in Developmental Cell Lineages

**DOI:** 10.1101/062539

**Authors:** Bradly Alicea, Thomas E. Portegys, Richard Gordon

## Abstract

Lineage trees of embryonic development contain much subtle information about the embryogenetic process. One type of information is contained in how the order of nodes are sorted at each level of the tree. Sorting of lineage trees is accomplished using a specific criterion for all levels (each representing a division event) of the tree, resulting in new types of trees (e.g. differentiation tree). Another type of information can be revealed from pairwise comparisons of each type of tree. By using a binary classifier to quantify the first type of information (ordering by level), we can obtain a quantitative measure for the second type of information by using the Hamming distance between equivalent positions in two trees. In this paper, we will introduce a method for calculating and visualizing the information content of embryogenesis called the information isometry technique. Information is extracted from developmental lineages using a binary classification system, and visualization is accomplished through the construction of isometric graphs, which re-represent a tree topology as a series of isometric lines. As the points representing each segment of an isometric line changes color, there is a shift in the underlying tree and its constituent cells. Isometric graphs reveal a number of patterns within cell lineages, including the relative information content of specific subtrees.

## INTRODUCTION

The identification of cells and all of their descendants during embryonic development involves constructing a lineage tree [1]. Invariant lineage trees of mosaic organisms such as invertebrates (*Caenorhabditis elegans*) [2, 3, 4], chordates (*Ciona intestinalis*) [5], and other animal species are highly predictable from organism to organism. This high degree of regularity enables higher-level analysis individual cells and their corresponding levels (each representing a division event) of the tree topology. The lineage tree representation of embryogenesis provides us with a directed acyclic graph (DAG) representation of cellular differentiation at multiple scales of complexity [6]. The horizontal ordering of cells in a lineage tree at each level generally represents their position on an anatomical axis. This order can be rearranged into a differentiation tree by using cell size criteria to determine the order of cells from left-to-right at each level.

We can use a binary code to classify each cell produced during a binary differentiation event (see Figure 1). This is a compact representation of cellular differentiation relative to both axial position or cell size variation. For both the lineage tree (axial position) and differentiation tree (cell size), we get a unique binary classification of each cell. Since a binary code allows for a direct comparison amongst different ordering criteria, we can ask whether the binary code is the same for a given cell across orderings, and whether that has any biological significance.

Using a binary classification system is not simply done out of computational expediency. The analytical framework presented here draws from work on the differentiation code [7, 8, 9] that demonstrates a means to classify cells and tissues as being part of a larger framework of pattern formation based on the dynamics of cellular differentiation. Differentiation codes are a generative modeling technique that results in a binary tree structure. To accomplish this, we order all descendent cells or tissue by a size criterion. We characterize each binary differentiation event as a comparison between smaller and larger units. In a binary tree rooted at the top of a two-dimensional graph (see Figure 1), the smaller units branch to the right, and the larger units branch to the left. Theoretically speaking, these branches correspond to contraction differentiation waves and expansion differentiation waves, respectively [7, 9]. In constructing an isometric graph, we generate both a differentiation code and a lineage code. The lineage code follows the same logic as the differentiation code, but is used merely to recapitulate the lineage tree order.

## SOURCE DATA AND METHODS

### Description of Dataset

This representative nematode (roundworm, Nematoda) is a usually hermaphroditic organism that has 959 cells in the adult hermaphrodite and 1031 in the adult male. Our secondary dataset [3, 4] consists of pre-hatching cell divisions (up to 558 cells). In the original acquisition of the data, identification of the cell nucleus was accomplished using a GFP+ marker.

Embryos for *C. elegans* embryos are incubated at 25^o^C [8], while the cell nucleus is imaged using a GFP+ marker. These techniques allow for approximation of volume as calculation of a sphere based on an approximation of cell diameter. Diameter, and nomenclature (identity) for all cells are extracted from observations of 261 embryos. These variables are then averaged over every observation of a specific cell type. Discrete GFP+ regions are segmented from microscopy images using computer vision techniques, and are used to construct an optimal spherical representation (see [3, 4]). Due to the nature of the quantification of the data, corresponding volumetric data of the whole embryo is not available. This spherical representation is based on identification of the centroid and annulus for these segmented regions from florescence signal intensity. While this is not an exact measure of diameter for the entire cell, it provides a reasonable approximation.

In order to address inconsistencies in size differences at the 2- and 4-cell stage of our secondary dataset, we generated a composite differentiation code. This combined knowledge of relative cell size at the 2- and 4-cell stage from cell polarity studies (see [10, 11]) with our dataset of cell volumes estimated from GFP+ nuclei. This composite differentiation code was used to estimate the *C. elegans* differentiation code. All raw data used in this study are available at Github. Lineage Tree Raw Data: https://github.com/balicea/DevoWorm/tree/master/Lineage%20Tree%20DB, Differentiation Tree Raw Data: https://github.com/balicea/DevoWorm/tree/master/Differentiation%20Tree%20Dataset. All processed data (used in the analyses) are available from the Open Science Framework: https://osf.io/pnh86/

### Differentiation Code and Tree Classification

Differentiation codes were constructed by using the binary differentiation events as described by Sulston et.al [2]. To arrive at a differentiation code, each daughter cell in a pair is classified as either “0” or “1”, dependent upon each cell’s relative volume. This classification is done independently of Eq. 1 and 2. The differentiation code is a sequence of binary digits that provide an absolute address in terms of lateral position (left-right) and depth (number of branches from the root) in the differentiation tree ordering. Lineage trees are coded according to their published arrangement in the lineage tree. Cells organized to the left are classified using a “0”, while cells organized to the right are classified using a “1”. The address system is the same as that of the differentiation tree, with the exception that the “lineage code” is in a different order with respect to the differentiation tree ordering. The distance between lineage and differentiation codes are calculated to arrive at a Hamming distance.

### Hamming Distance Calculation

The Hamming distance metric [12] is used to calculate a distance between differentiation or lineage codes for two cells with the same depth (see Figure 1, panel D). The Hamming distance is defined as

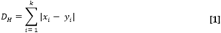

where *x_i_* and *y_i_* are binary strings of the same length (depth in the tree) compared pairwise, k is the length of both strings, and DH counts the number of differences between the two strings in terms of bits. This calculation can be done for the same cell, in regards to its positions in the lineage and differentiation trees, between different cells in the same tree, that are at the same depth, or between cells at the same depth in trees of two different organisms.

In this paper, we calculate the Hamming distance between the orderings provided by the lineage tree representation versus the differentiation tree representation. This is done to provide a quantitative measure of how ordering the line of cellular descent by different criteria changes the relationships between these trees. In the random example, each comparison is made on a unique hypothetical cell that represents part of an ordered set. These data are then reordered (randomized) using the MATLAB function shuffle.m. In the *C. elegans* example, each comparison is made on a unique biological cell as defined by the Sulston et.al [2] nomenclature. A unique cell will always have the same depth, but a different binary code of the same length, which is dependent upon the tree ordering.

## RESULTS

To evaluate this, we conduct a quantitative comparison of equivalent graph topologies by classifying the nodes of a tree structure using information theory. More specifically, we measured the Hamming distance between each pairwise comparison of the same cell. To accomplish this, the tree structure is rooted in one order and compared with another order. This results in an isometric graph that reveals the information isometry between embryogenetic trees sorted and classified by different criteria in a bivariate manner.

An isometric graph is equivalent to a lineage tree without branches rotated clockwise 45 degrees. Isometric graphs are organized in the following way: each cell is plotted as a point in bivariate space. The diagonal line on an isometric graph is represented by a line of points that intersects both the *x* and *y* axes at identical values. The isometric graph in Figure 2 demonstrates how x and y abscissa both occur at intervals of 1, 3, 7, 15, and 31. The *x* and *y* axes represent the right-hand and left-hand sides of the cell lineage tree, respectively.

To demonstrate how this works, we have created a cell lineage and their corresponding lineage and differentiation codes using pseudo-data (Supplemental File 1). We can use this as a proof-on-concept before moving on to empirical examples. First, we generated a lineage tree topology and then resorted the order of these cells randomly at each level. The comparison presented in an isometric graph is shown in Figure 2. As a generality in the random case, subsequent differentiation events lead to progressively larger Hamming distances. A Hamming distance of 3 seems to dominate the comparison between tree orderings after the fourth level, but this seems to be due to the effects of averaging. Across the five replicates, the Hamming distance fluctuates amongst the first four levels, but converge upon a monotonically increasing function afterwards as shown in Table 1.

**Figure 1.**
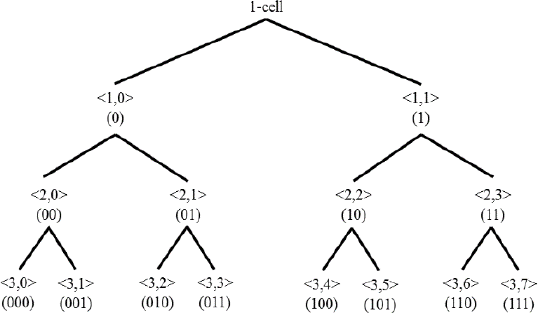
Demonstration of lineage/differentiation code and how it maps to a two-dimensional position in the tree topology. Numbers in brackets represent the ordered position of each node (cell), while numbers in parentheses represent the binary code generated from the order and level of a given node (cell).

**Figure 2.**
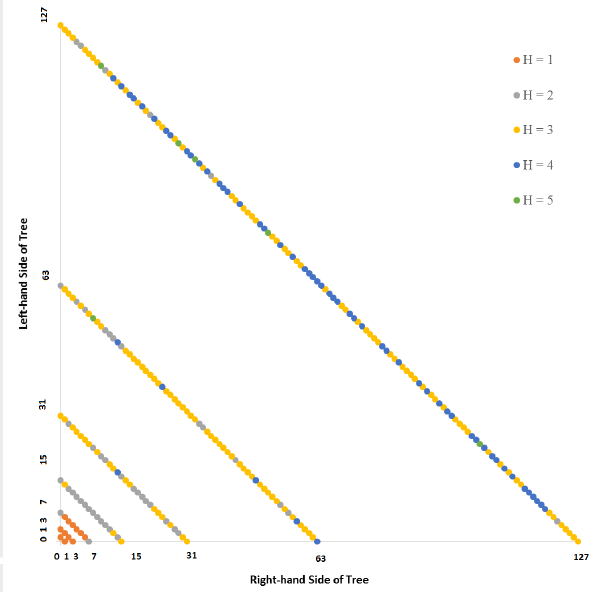
Isometric graph based on the averages for five randomized lineage trees, seven (7) levels apiece relative to a lineage code order. The *H* abbreviation stands for Hamming distance. The position of a point representing a cell is based on the depth of its node in the differentiation tree. The positions of all points are rotated 45 degrees clockwise from a bottom-to-top differentiation tree ordering (where the 1-cell stage is at the bottom of the graph). Each cell is colored with its Hamming distance. Along a given line, the cells appear in their order, left to right, in the differentiation tree.

**Table 1.** The average Hamming distance and standard deviation per tree level for both the random case and *C. elegans* biological example. SD = standard deviation of the mean.

| Level | Random Mean | Random SD | *C. elegans* Mean | *C. elegans* SD |
| --- | --- | --- | --- | --- |
| 1 | 0.6 | 0.5 | 1.0 | 0.0 |
| 2 | 1.2 | 0.7 | 2.0 | 0.0 |
| 3 | 1.2 | 0.9 | 2.5 | 0.5 |
| 4 | 2.0 | 0.9 | 3.1 | 0.6 |
| 5 | 2.6 | 1.2 | 3.9 | 0.7 |
| 6 | 3.0 | 1.1 | 4.4 | 1.1 |
| 7 | 3.4 | 1.4 | 4.5 | 1.1 |

The *Caenorhabditis elegans* isometric graph is based on the lineage tree *sensu* Sulston [2], and is shown in Figure 3, the source data for which can be found in Supplementary File 2. In this case, the lineage tree is compared to the differentiation tree. Recall that unlike the generation of random orderings at each level of the lineage tree, the lineage tree and differentiation tree are both invariant across organisms. In this case, the average Hamming distance and standard deviation increases monotonically with level as shown in Table 1.

**Figure 3.**
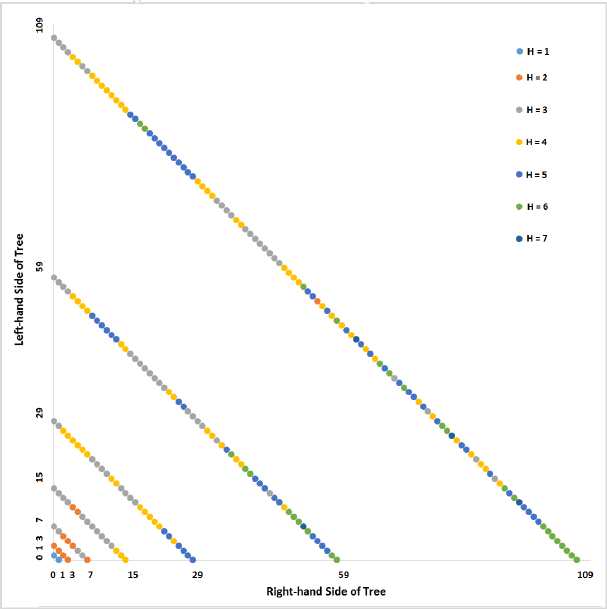
Isometric graph based on seven (7) levels of the *Caenorhabditis elegans* lineage tree relative to a lineage code order.

A comparison of the random case and *C. elegans* is shown in Figure 4, and illustrates what we should expect from an orderly process versus a randomized process. In this graph, we compare the mean Hamming distance for each level of isometric graphs resulting from the random case (Figure 2) and *C. elegans* (Figure 3). Early cell polarity patterns in the 2‐ and 4-cell stages of the *C. elegans* embryo [9] result in a higher average Hamming distance between the lineage and differentiation codes as compared to the random condition. When cell polarity is not taken into account, the trend across levels for *C. elegans* is consistent with the random case (albeit more linear). The main difference between the two conditions involves a monotonically-increasing mean value for the Hamming distance as cellular differentiation proceeds from the 1-cell case.

**Figure 4.**
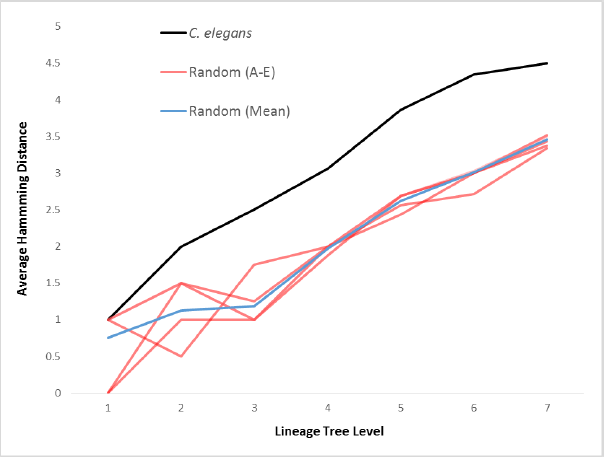
Plot of Hamming distance for six averaged series: five randomized orders for each level of (red, based on pseudo-data) and *C. elegans* (black, based on lineage tree *sensu* Sulston).

By contrast, the random case shows wide variation for the early stages of differentiation, likely due to the random positioning of a relatively small number of cells. By the third to fifth level, the randomly-generated Hamming distances begin to converge to monotonically-increasing functions. This is an averaging effect on an increasingly larger number of cells, but is contingent upon earlier stages of embryogenesis. In fact, we observe a slight divergence between the averaged random case and the *C. elegans* case (which itself is based on an ensemble average of 261 embryos).

Taken together, Figures 2, 3, and 4 also show that although subtree-specific changes in the Hamming distance can be consistent with both the biological and random versions of an isometric graph, differences between the lineage and differentiation codes tend to increase as development unfolds. In other words, comparing the differentiation and lineage codes reveals an aggregate statistical signal that may be biologically meaningful.

To find out what this biological significance might involve, we examined Hamming distance by subtrees rooted at the 8-cell stage. This provided us with eight categories. For each category, a histogram of Hamming distances were generated and plotted in Figures 5 (random) and 6 (*C. elegans*). In Figure 5, we establish the variation in Hamming distance distributions that exist for the averaged randomized case. In this case, even though there is some skew in the eight distributions, we also observe a degree of uniformity defined by a Hamming distance of three. Examining the pseudo-data for all five randomized tree orderings reveals a similar pattern, albeit with more variance (see Supplemental File 1).

**Figure 5.**
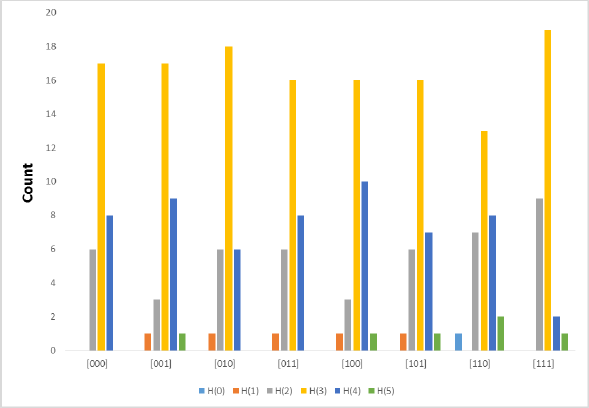
Histogram of subtrees rooted at the 8-cell embryo based on averages for five randomized linage trees, seven (7) levels apiece.

**Figure 6.**
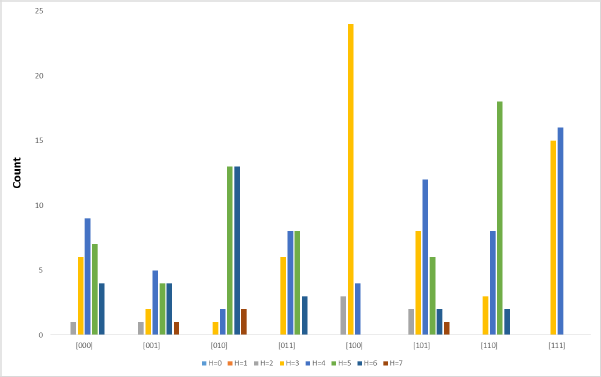
Histogram of subtrees rooted at the 8-cell embryo for seven (7) levels of the *Caenorhabditis elegans* lineage tree.

Figure 6 demonstrates a biologically-meaningful signal and a difference in Hamming distance distributions. In this case, two sublineages (“100” and “111”, descendents of the AB founder cell) demonstrate an extreme bias for a Hamming distance ranging from 2-4 for sublineage “100” and 3-4 in sublineage “111”. This lack of variation is an indicator that these sublineages preserve their order between the lineage tree and differentiation tree.

In the *C. elegans* case, we can further examine this signal by using partial isometric graph. These are shown in Figure 7 and 8. In a partial isometric graph, only selected sublineages are displayed, but are displayed to their full depth (based on available data). Figure 7 shows Hamming distance data for subtrees “100” and “111”. Indeed, we can observe evidence of this conservation by sublineage, as there are blocks of two or more cells in a row with the same Hamming distance rather than a heterogeneous pattern (more information content). This is particularly true of the later-occurring levels of the trees. In Figure 8, the “000” and “001” subtrees show more of a heterogeneous pattern of Hamming distances.

**Figure 7.**
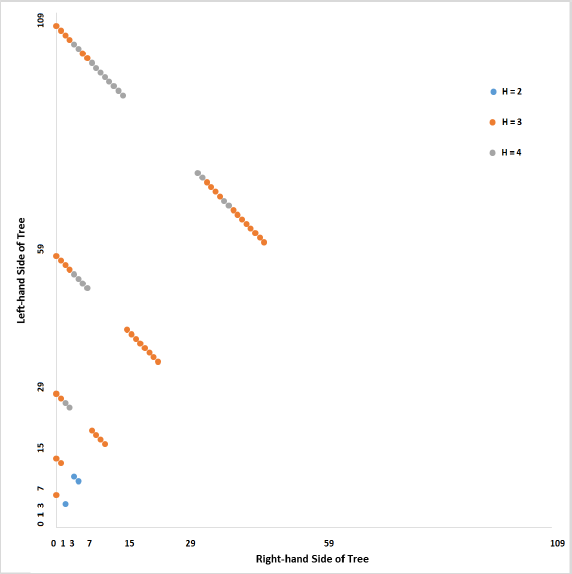
Isometric graph based on seven (7) levels of the *Caenorhabditis elegans* lineage tree (sampled to include only the “100” and “111” sublineages) relative to a lineage code order.

**Figure 8.**
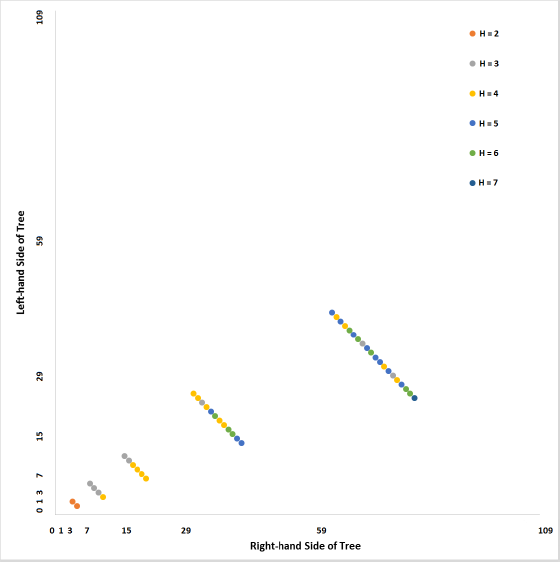
Isometric graph based on seven (7) levels of the *Caenorhabditis elegans* lineage tree (sampled to include only the “000” and “001” sublineages) relative to a lineage code order.

Taken together with the histogram data shown in Figure 6, we can surmise that the lineage to differentiation code comparison results in a weak biological signal. While we can distinguish between the biological case (*C. elegans* developmental cell lineage) and the random case, we are still unsure as to the differences between biological signal and statistical noise. While this is hard to extract at the aggregate level or in terms of statistical distributions, isometric graphs allow us to acquire additional information about these relationships and relate them directly to the biology of embryogenesis.

## DISCUSSION

Application of an information theoretic approach to investigation of lineage trees and embryogenesis more generally provides a number of advantages. Information theory and visualization allows us to apply a rigorous quantitative approach to understanding patterns of cellular differentiation at multiple spatial and temporal scales. The ability to compare different organizations attributes of the embryogenetic process within a common framework provides a flexible and potentially universal representational framework. This may give a basis for investigating within and between species variation. Overall, the coupling of differentiation/lineage codes with the isometric graph approach provides a parsimonious approach to finding patterns within and between cell lineage subtrees.

There also appears to be a difference between a biological signal and random noise. Random noise seems to be defined by either a heterogeneous pattern along a single isometric line (Supplemental File 3), or convergence to a Hamming distance of 3 after averaging (Figure 2). This is systematically lower than in the case of *C. elegans*, where the combined effects of early cell polarity and the founding of functionally distinct subtrees results in a more informative differentiation code.

By contrast, a biological pattern can be defined by local conservation (less information content) within a sublineage, and appears as a series of two or more consecutive points with the same Hamming distance. This can be observed by looking at the coordination of daughter cell criteria over several differentiation events. When the Hamming distance of pairs of daughter cells changes together over developmental time, it serves as a signal of this coordination [8, 13]. While we have focused on the comparison between differentiation and lineage codes in this paper, binary classification codes can be model for other criterion, and thus allowing for multiple dimensions of variation to be explored one pairwise comparison at a time.

This leads us to ask what is biologically unique regarding the cells in the “100”/”111” sublineages versus the “000”/”001” sublineages of the *C. elegans* embryogenetic trees. The fact that there are observable differences between sublineages for the biological example demonstrates that this method provides a signal distinct from random noise. The “100”/”111” sublineages are part of the AB sublineage, the cells of which tend to exhibit a generalized fate. By contrast, the “000”/”001” sublineages (sublineages C, D, and P) are overrepresented by cells that contribute to germ line formation and muscle cells [13]. Each sublinage also possess additional characteristics such as their own internal cell cycle clock [14]. While an information-theoretic approach does not pinpoint these causal factors directly, it does provide a means for identifying features for further investigation.

Finally, we would like to revisit Figure 3 (entire *C. elegans* developmental cell lineage up to 7 division events) and inquire about the identities of cells with a Hamming distance of 6 and 7. As Figure 6 and 7 demonstrate, sublineages such as C and D feature cells with a Hamming distance of 6 and 7, while other sublineages may not feature any such cells. To systematically investigate this, all cells with a Hamming distance of 6 and 7 were identified and classified into their corresponding developmental sublineage (AB, C, D, E, or MS). This allows for the proportion of cells with a Hamming distance of 6 and 7 in each sublineage to all cells at levels 6 and 7 in a given sublineage to be calculated (Table 2). Upon comparison, we can see that a plurality of cells in the D sublineage and a majority of cells in the MS sublineage fit these criteria. What this means in terms of terminal differentiation is not clear. While all cells in the D sublinege contribute to muscle formation, the MS sublineage is a bit more diverse, with cells contributing to the formation of muscle, neuron, glands, and coelomocytes [2]. As in the case of Figure 7, these sublineages are located towards the posterior end of the nematode.

**Table 2.** Identity of cells with Hamming distances of 6 and 7 by sublineage, the number of such cells by *C. elegans* developmental sublineage, and the proportion of these counts by the total number of cells at levels 6 and 7 of the *C. elegans* lineage tree.

| Sublineage | Number of Cells | Proportion |
| --- | --- | --- |
| AB | 5 | 0.052083333 |
| C | 4 | 0.166666667 |
| D | 5 | 0.416666667 |
| E | 3 | 0.125 |
| MS | 16 | 0.666666667 |

## ACKNOWLEDGEMENTS

We would like to thank Drs. John E. Sulston and Zhirong Bao for access to nomenclature and biometric data for the *C. elegans* embryo. We would also like to thank Dr. Stephen Larson and the OpenWorm Foundation for their institutional support, and George E. Mikhailovsky for his comments and suggestions.

